# Unisensory visual and auditory objects are processed in olfactory cortex, independently of odor association

**DOI:** 10.1101/2023.04.20.537709

**Authors:** Evelina Thunell, Moa Peter, Behzad Iravani, Danja K. Porada, Katharina Prenner, Fahimeh Darki, Johan N. Lundström

## Abstract

Primary sensory cortices have been demonstrated to process sensory input from non-preferred sensory modalities, e.g. primary visual cortex reacting to auditory stimulation, bringing their presumed sensory specificity into question. Whether this reflects processing of the non-preferred stimulus per se or originates from cross-modal associations is debated. Visual/auditory objects typically have strong reciprocal associations; hence, it is difficult to address this question in these modalities. Here, we attempt to dissociate between the two competing hypotheses of whether this form of activation in primary cortices is caused by unisensory processing or cross-modal associations by turning to the olfactory system where cross-modal associations are generally weaker. Using unisensory visual and auditory objects with odor associations ranging from none to strong, we show that the posterior piriform cortex, an area known to process odor objects, is activated by both sounds and pictures of objects. Critically, this activation is independent of the objects’ odor associations, thereby demonstrating that the activity is not due to cross-modal associations. Using a Floyd–Warshall algorithm, we further show that the amygdala mediate condition-relevant information between the posterior piriform cortex and both the auditory and visual object-oriented cortices. Importantly, we replicate past findings of clear crossmodal processing in the visual and auditory systems. Our study demonstrates processing of non-olfactory input in olfactory cortices that is independent of cross-modal associations and contributes to a more nuanced view of modality specificity in olfactory, auditory, and visual cortices.

## Introduction

The concept of a unisensory cortex has been challenged by demonstrations that primary sensory cortices in healthy individuals sometimes respond to unisensory stimuli of a different sensory modality. For example, the primary visual cortex (V1) is commonly activated by visual stimulation but, under certain circumstances, also responds to auditory and tactile stimulation: e.g in humans by abstract spoken words [Seydell-Greenwald et al., 2020] and, in rats, during tactile exploration [Vasconcelos et al., 2011]. Two competing explanations have emerged for why a primary cortical area responds to sensory stimuli from other modalities: 1) activations are due to *cross-modal associations*, i.e. association of a non-preferred sensory stimulus to a preferred sensory stimulus, or 2) alternatively, activations constitute *true unisensory processing of non-preferred sensory stimuli*, meaning that the stimulus of the non-preferred sensory modality is itself processed in the primary sensory area. Dissociating these theories in the audio-visual domain is a non-trivial task given the tight ecological connection between the visual and auditory systems where e.g. spoken words (whether real or abstract) and the lips of the person uttering them are inherently associated. The olfactory system does not come with the same caveat. Odors are difficult to identify without other cues, odor sensations often lack spontaneous associations to other modalities, and objects that do not contain organic materials, such as a clock, are seldom associated with odors [Cain, 1979]. In the olfactory system, it is thus possible to dissociate the two competing theories of why a main sensory cortical area responds to stimuli of non-preferred sensory modalities.

The main olfactory cortex has long been considered to comprise the piriform cortex due to its role as the primary recipient of direct projections from the olfactory bulb [Mainland et al., 2014; Zald and Pardo, 2000]. Similarly to activation by non-preferred stimuli in our other sensory cortices, there are several accounts of the piriform cortex processing both unisensory non-olfactory stimuli [Gottfried et al., 2004; Maier et al., 2012; Porada et al., 2019; Zhou et al., 2019] and integrating multisensory information [Lundström et al., 2019; Porada et al., 2019]. Indeed, direct anatomical connections to piriform cortex from the retina [Cooper et al., 1994], auditory system [Budinger and Scheich, 2009; Budinger et al., 2000], and gustatory system [Datiche et al., 1996; Johnson et al., 2000] have been found in several non-human mammals and provide a potential route for sensory information from other sensory modalities into initial processing stages of the olfactory system. As with the other sensory cortices, it is unclear whether cross-modal associations or true unisensory processing of non-preferred stimuli drives the activation [Ohla et al., 2017]. In most studies that demonstrate piriform cortex activation by non-olfactory unisensory stimuli, the stimuli were associated with odors either through presentation of odors during the experiment or by ecologically or experimentally induced multisensory associations [Olofsson et al., 2019; e.g., Schulze et al., 2017]. Non-preferred stimulus processing in the piriform cortex extends beyond low-level sensory information to associative processes, such as odor memory [Yang et al., 2021], attention [Veldhuizen and Small, 2011; Zelano et al., 2005], and cross-modal processing [Gottfried et al., 2004; Lundström et al., 2019; Zhou et al., 2019].

To dissociate between the two competing hypotheses, “cross-modal association” and the “true unisensory processing of non-preferred stimuli”, we presented unisensory visual and auditory objects with varying degree of odor associations, from very weak (e.g., a church bell) to strong (e.g., popcorn), to participants while assessing neural responses with functional magnetic resonance imaging (fMRI). Critically, to keep participants agnostic to olfaction and remove any nuisance intrinsic modulation, the word “odors” and related terms were not mentioned until after the scanning session when participants rated how much they associated each visual and auditory object to an odor. This design allowed us to dissociate between the two competing hypotheses because if potential activity in piriform cortex is dependent on cross-modal association, the response to an object should reflect how much the individual associates it to an odor. On the other hand, if activity in piriform cortex is due to true unisensory processing of non-olfactory stimuli, the neural response should be independent of odor association.

The anatomical separation between the anterior and posterior piriform cortex has been suggested to correspond to a functional specialization where the anterior portion (APC) processes mainly low-level olfactory features whereas the posterior portion (PPC) is more involved in odor object processing [Gottfried, 2010; Lundström et al., 2011]. This separation is analogous to the hierarchical organization of the visual and auditory systems where activity in the primary visual cortex (V1) and primary auditory cortex (A1) reflects mainly low-level features whereas the lateral occipital complex (LOC) and higher order auditory cortex (hAC) process objects [Porada et al., 2019]. Here, we used fMRI to assess potential processing of non-preferred stimuli in both low- and high-level sensory cortices with both region of interest (ROI) and whole-brain analyses.

## Methods

Stimuli, motion parameters, and imaging results are available on the open science framework (OSF) repository: https://osf.io/duwrb/?view_only=d214b458aee64204bf6e09df38249631.

### Participants

A total of 54 individuals participated in the experiment. Data from seven of these were excluded due to excessive sleepiness as defined by large amount of eye closure, judged from eye-tracker monitoring, or excessive head movement detected during pre-processing of the data. The remaining 47 individuals (28 women; mean age 33.5 years, *SD* = 10.1, range 18-46 years) reported being healthy and having normal or corrected-to-normal visual acuity, normal hearing (except for 1 participant who had reduced hearing on one ear), and normal olfactory function. The procedures were in accordance with the Declaration of Helsinki and approved by the Swedish Ethical Review Authority (Dnr. 2020-06533). All participants provided written informed consent prior to the experiment.

### MR and physiological data acquisition

Neural responses to visual and auditory stimuli were assessed using a 3 T Magnetom Prisma Scanner (Siemens) with a 20-channel head coil. We acquired functional data using a 3D echo planar (EP) sequence (56 slices, repetition time (TR) = 1700 ms, echo time (TE) = 30 ms, flip angle (FA) = 70°, voxel size = 2.234 mm x 2.234 mm x 2.200 mm, field of view (FoV) = 94 x 94 voxels). Slices were tilted 20 degrees. A structural image was also obtained using a 3D MPRAGE T1-weighted sequence (208 slices, TR = 2300 ms, TE = 2.89 ms, FA = 9°, voxel size = 1 mm x 1 mm x 1 mm, FoV = 256 x 256 voxels). At the end, we acquired a narrow T2-weighted image that was not used in this study. Participants were instructed to breathe through their nose throughout the experiment and their respiratory trace was continuously recorded during the functional runs using birhinal nasal cannulas attached to a PowerLab 4/26 unit via a spirometer (ADInstruments Inc., Colorado Springs CO, USA). To evaluate participants’ wakefulness during the functional runs, we continuously monitored their eyes using a TRACKPixx3 2kHz binocular distance eye tracker (VPixx Technologies Inc., Saint-Bruno, QC Canada).

### Stimuli and task

The stimulus set comprised grayscale pictures and sounds of 48 easily identifiable objects, chosen to evoke various degrees of odor association. We selected this set based on a pilot study where 23 healthy participants identified and rated the level of their odor association to the pictures and sounds of 60 everyday objects (See Supplementary material Section 1 and Figure S1 for details). Sounds were obtained from the BBC Sounds Database, http://bbcsfx.acropolis.org.uk, and Freesound, https://freesound.org, and pictures were taken from the Open Image Dataset, https://opensource.google, and ImageNet, http://www.image-net.org. The sounds were seamlessly looped or trimmed to last 3 seconds and equalized with a root mean square equalization to -23 dB loudness [Regenbogen et al., 2016]. Pictures were cropped symmetrically from all sides to square shape with a side length equal to the shortest of the original height and width and converted to grayscale using the rgb2gray function in Matlab. The visual stimulation was presented on a 40’’ 4K ultra-HD LCD display (NordicNeuroLab AS, Bergen, Norway) placed behind the scanner bore and viewed via a head coil mirror. The viewing distance was approximately 180 cm, and the pictures spanned 4.5 degrees of visual angle. The auditory stimulation was presented via binaural headphones (NordicNeuroLab AS, Bergen, Norway), at a loud volume that was adjusted when needed to be tolerable to the participant. All objects were always presented to participants as unisensory stimuli to avoid any additional crossmodal training beyond the inherent associations between a sound and image portraying the same object.

### Procedure and design

#### Practice

To ensure recognition of the objects, all stimuli were presented prior to MR scanning one at a time together with the name of the object. During this pre-scanning practice, all pictures appeared one time with the object name written underneath, and all sounds were played one time with the object name written in the middle of the screen. Participants were instructed to learn to identify the objects and told that they would later be tested on this task. While there was no subsequent explicit behavioral test, they were asked to identify the objects during the fMRI scanning.

#### MR scanning

The MR session included four functional runs. Each run lasted 10 min and 15 s and contained eight blocks lasting 1 min each. Within a block, six stimuli were presented, each with a duration of 3 s and preceded by 7 s ± 3 s of no stimulus. All blocks were preceded and followed by 15 s of no stimulus. The stimuli were grouped into pure sound and pure picture blocks. Further, the stimuli were grouped according to the average odor association level across the two modalities in the pilot study, such that the stimuli in each block belonged to either the highest or lowest rated half. The low and high odor association blocks were interleaved, with half of the participants starting with a low and the other half starting with a high odor association block. All 96 stimuli (48 sounds and 48 pictures) were presented once in the first half of the runs (runs 1 and 2), and a second time in the second half of the runs (runs 3 and 4), in a random order. The participants’ task was to fixate a cross that was presented throughout each run except during picture presentation, and to identify each presented object without giving any response.

#### Odor association

After the MR session, participants rated how much they associate each stimulus with an odor using a visual analogue scale ranging from 0 (no odor association) to 100 (very strong odor association). (This question was posed in a general fashion and thus did not refer to the MR scanning in particular.) Pictures and sounds were presented in alternating blocks with 12 stimuli per block. Critically, this task was the first mention of odors to the participants and no odors were presented at any point.

### Imaging data processing and statistical analyses

Pre-processing was carried out separately for each participant using SPM12 (version 7487; Penny et al., 2011), including slice time correction, motion correction, coregistration of the functional runs, coregistration of the anatomical and functional data, and normalization to MNI space. The functional data were smoothed for whole-brain analyses using a 6 mm full width at half maximum kernel, while region of interest (ROI) analysis was performed on non-smoothed data. For one participant, the first functional run was excluded due to a headphone problem which was remedied before the three remaining runs.

First level statistical analyses were conducted using a General Linear Model (GLM), as implemented in SPM12. The BOLD signal responses to the pictures and sounds were modeled separately as a boxcar convolved with a canonical hemodynamic response function (HRF) as basis function. We used each participant’s individual odor association ratings of the sounds and pictures, which were collected *after* the MR data acquisition, as parametric modulators (orthogonalized). To account for potential confounding motion effects, we added the six rigid-body motion parameters (i.e., 3 translational and 3 rotational variables) from the motion correction as regressors of no interest. The four regressors of interest were response to pictures, parametric modulation of the response to pictures by odor association, response to sounds, and parametric modulation of the response to sounds by odor association. We computed simple t-contrasts of each of these against implicit baseline.

Second level whole-brain analyses were conducted using one-sample t-tests on the four regressors of interest (participants’ responses to pictures, modulation of these responses by the odor association ratings of the pictures, responses to the sounds, and modulation of these responses by the odor association ratings of sounds). In a ROI analysis, we extracted mean contrast estimates in six bilateral ROIs representing basic and object-related neural processing within each sense. Namely, anterior piriform cortex (APC) and posterior piriform cortex (PPC) for olfaction, primary visual cortex (V1) and lateral occipital complex (LOC) for vision, and primary auditory cortex (A1) and higher order auditory cortex (hAC) for audition. These ROIs (Porada et al. 2019) are based on a combination of functional activations in previous studies and standardized anatomical masks to maximize specificity and independence. For each participant and ROI, we extracted mean contrast averages from the left and right hemispheres separately and subsequently averaged over the two hemispheres. We performed one-sample t-tests on the extracted voxel-averages for each ROI and contrast using an α-level of .05. Whole-brain activations for these four effects are presented corrected for multiple comparisons using a peak-level threshold at *p* < .05 family-wise error (FWE). The imaging data had acceptable coverage in all ROIs (participant averages: APC: 76%±12, PPC: 64%±13, V1: 98%±2, LOC: 89%±6, A1: 98%±2, hAC: 82%±8).

In an additional analysis, the ROI analysis described above was repeated with all conditions split up in low and high odor association groups based on the participants’ individual ratings, separately for pictures and sounds. The 4 regressors of interest were thus replaced by 8 regressors (response to pictures with weak and strong odor association, parametric modulation by odor association of the response to weak and strong odor association pictures, response to sounds with weak and strong odor association, and parametric modulation by odor association of the response to sounds with weak and strong odor association). We also performed an additional analysis where each stimulus was assigned a separate regressor. In this GLM, there were in total 96 regressors of interest (48 for pictures and 48 for sounds); no parametric modulation was added. We ranked the stimuli according to each individual’s smell association ratings, with random ordering in case of identical ratings for more than one stimulus, and correlated these rankings with the ROI activations, separately for pictures and sounds.

### Dynamic functional connectivity and path analyses of signal from the visual/auditory cortices to the piriform cortex

To

How the visual and auditory cortices communicate information to the olfactory cortex is not known. We therefore performed an analysis of most direct path by extracting BOLD activities from 272 spherical ROIs across the whole brain from on the CONN functional connectivity toolbox atlas [Whitfield-Gabrieli and Nieto-Castanon, 2012] (radius = 18mm, Figure 5A). The BOLD signals were denoised through a standard denoising pipeline [Nieto-Castanon, 2020], including regression of confounding variables. Specifically, these variables encompassed motion parameters along with their first-order derivatives (12 factors) as well as adjustments for condition-related effects (2 factors: pictures and sounds). The denoised BOLD data were windowed to five-samples long segments using a Hanning sliding window with 50% overlap (alternating between 2 and 3 samples overlaps). Subsequently, we computed the Pearson correlation between the 272 brain ROIs within each window, yielding time-series data representative of the dynamic functional connectivity (dFC; Figure 5B).

On the individual level, we constructed a linear regression analysis to assess the condition-dependent modulation of the dFC (Figure 5C). A design matrix with intervals labeled either sound or picture was created. These intervals represent the consecutive time points where each stimulus was presented. To account for the hemodynamic response, the design matrix was convolved with a HRF. The linear regression determined the extent that the condition (i.e., sounds and pictures) affected the brain-wide dFCs, yielding individual connectivity beta values. At the group level, the beta values underwent a *t*-transformation and were averaged across cerebral hemispheres. To eliminate false positive connections, the *t*-transformed maps were thresholded based on two criteria: 1) any ROI with averaged gray matter probability falling below 40% was discarded; 2) only the top 20% of the strongest connections were retained (Figure 5D).

We opted to focus on LOC and hAC based on the assumption that the olfactory system preferentially integrates object-related information rather than the low-level features processed by the primary visual and auditory areas (V1 and A1). In general, the PPC receives more input from other cortical areas than does APC [Cohen et al., 2017], and we therefore consider only PPC in this analysis. To assess the functional connectivity between PPC and object processing areas within visual (LOC) and auditory (hAC) cortices, we converted the group *t*-transformed and thresholded dFC connectivity matrix to distance matrices, by inverse transform. The inverse transformation was required to be able to use Floyd–Warshall algorithms [Cf. Fransson et al., 2011] to find the shortest path between each pair of nodes, i.e., between auditory and visual object processing areas and the PPC (Figure 5E).

## RESULTS

### Olfactory cortex processes unisensory visual and auditory stimuli, independently of odor association

We first tested whether the PPC processes unisensory non-olfactory objects using ROI specific analysis given our directed hypothesis. We found a significant activation by pictures of objects in the PPC ROI (Figure 1A-B, *t*(46) = 4.96, *p* = 9.99*10^-6^, Cohen’s *d* = 0.72). Similarly, we found an activation by sounds of objects in PPC (Figure 1B, *t*(46) = 8.82, *p* = 1.89*10^-11^, Cohen’s *d* = 1.29). Next, we assessed whether the pictures and sounds also activated the APC. While we did find a significant activation in APC by sounds (Figure 1B, *t*(46) = 6.04, *p* = 2.51*10^-7^, Cohen’s *d* = 0.88), there was no apparent activation by pictures (Figure 1B, *t*(46) = 0.24, *p* = .81, Cohen’s *d* = 0.04).

**Figure 1.**
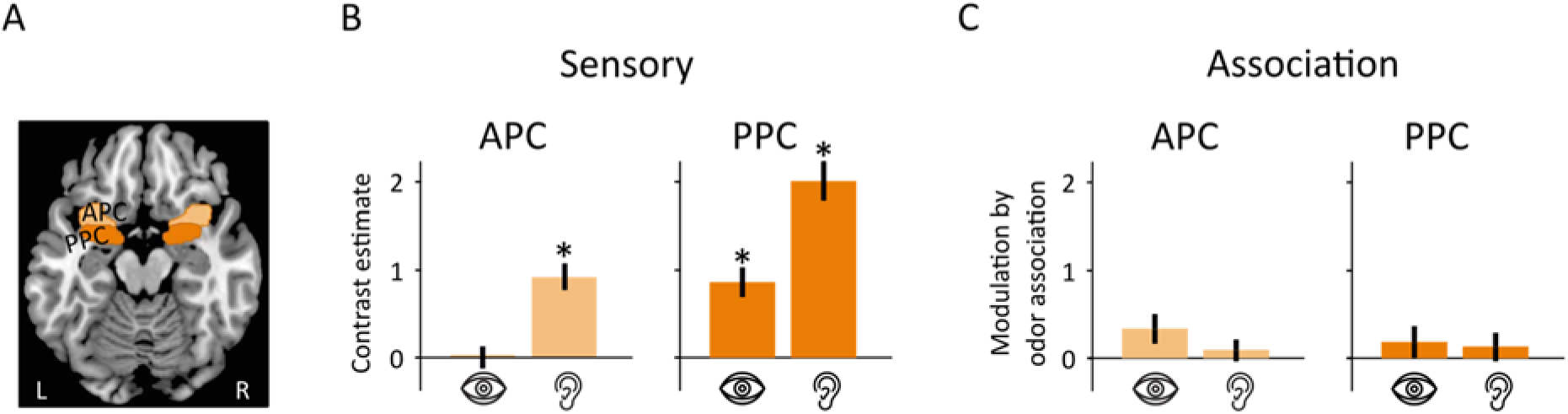
Region of interest MR imaging results for olfactory brain areas. **A)** Location of olfactory ROIs. **B)** In Anterior Piriform Cortex (APC), the extracted mean contrast estimates are significantly above zero for sounds (ear symbol) but not for pictures (eye symbol). In Posterior Piriform Cortex (PPC), both pictures and sounds elicited significant activations. **C)** In both APC and PPC, the mean parametric modulation by odor association is not significantly different from zero for sounds (ear symbol) nor for pictures (eye symbol). Asterisks mark statistical significance. Error bars depict ± 1 Standard Error of the Mean (SEM).

We used the ROI-based approach due to our region-specific hypothesis. A more conservative approach is a whole-brain analysis with correction for multiple comparisons across voxels. Using a whole-brain analysis, we could confirm the response to sounds in both PPC and APC using a conservative peak-level threshold of *p* < .05 FWE (Figure 2 A-B). The PPC response to pictures is not significant at this level but is visible with a more liberal uncorrected criterion of *p* < 0.001 (Figure 2B). In APC, no voxels were significantly activated by pictures, even with the liberal uncorrected criterion (Figure 2A).

**Figure 2.**
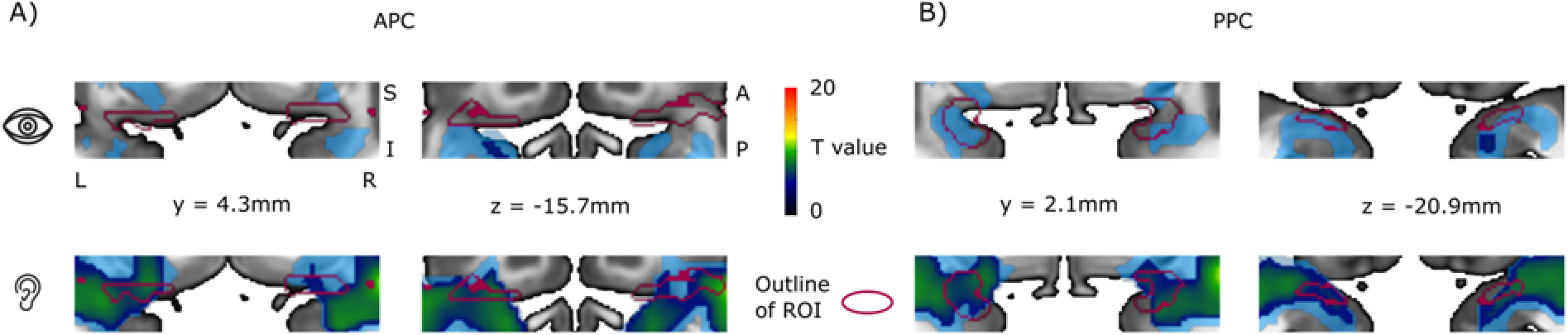
Whole-brain MR imaging results. A) APC. Activation by pictures (eye symbol) and sounds (ear symbol) are overlaid on a transverse and coronal cut-out covering the APC region of interest. Confirming the ROI results, the activation by sounds, but not that by pictures, is significant. Color bar indicates significance at the *p* < .05 family-wise error (FWE) level across the whole field of view. The semi-transparent light blue indicates significance at the *p* < 0.001 level, uncorrected for multiple comparisons. B) PPC. The activation by sounds, but not pictures, is significant in the PPC at the *p* < .05 FWE level.

These results demonstrate that olfactory cortex responds to unisensory visual and auditory objects. Critically, the activations occurred in the absence of any olfactory stimulation or explicit odor associations. No odors were presented at any point during the experiment and odors were first mentioned to the participants during the post-scanning association task. To test whether the activations were driven by potential odor associations that might have been automatically evoked when an object was viewed or heard, we assessed the neural modulation by participants’ degree of odor association to each object. Importantly, these ratings were obtained post-scanning and therefore could not influence the neural activation (Figure 3).

**Figure 3.**
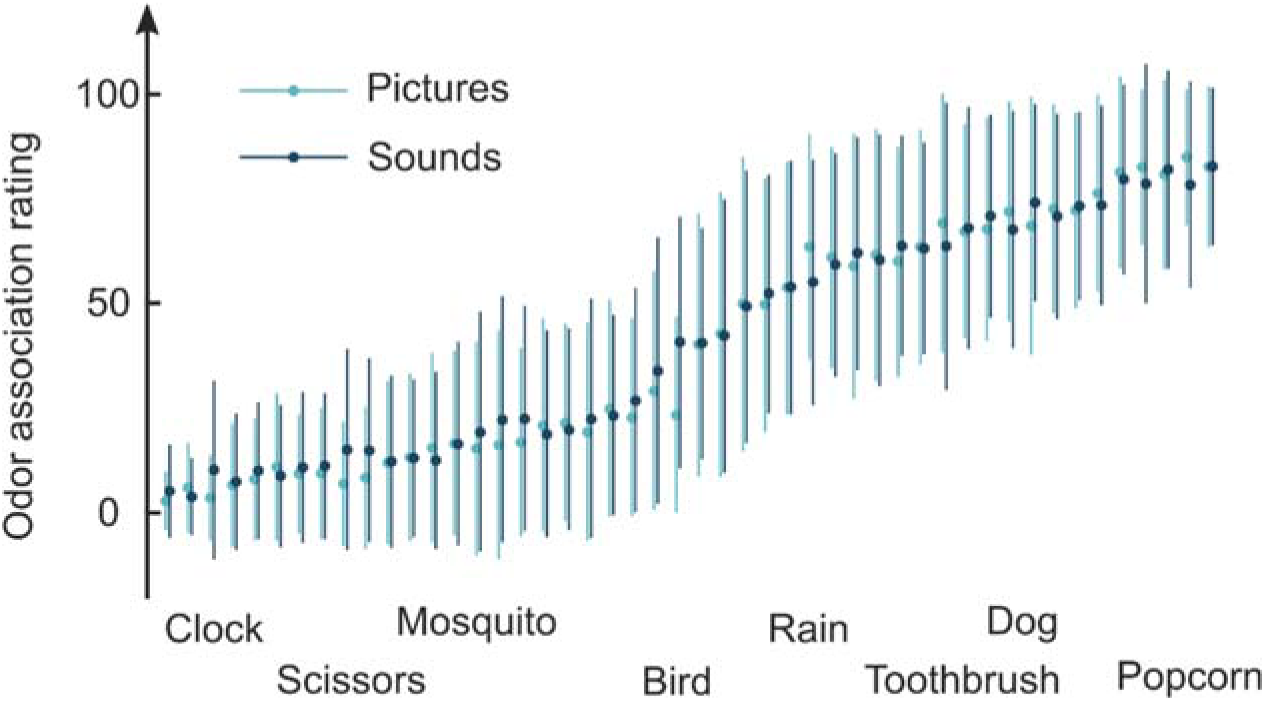
Average odor association ratings for the 48 objects. Overall, the picture and sound representation of the same object have similar odor association. Critically, values span a large portion of the scale. Eight example objects are named below the graph. Error bars denote ±1 Standard Deviation (SD).

There was no significant parametric modulation by odor association in the PPC for either pictures (Figure 1C, *t*(46) = 0.96, *p* = .34, Cohen’s *d* = 0.14) or sounds (Figure 1C, *t*(46) = 0.75, *p* = .46, Cohen’s *d* = 0.11). Likewise, the modulation by odor association of the response to sounds in the APC was non-significant (Figure 1C, *t*(46) = 0.65, *p* = .52, Cohen’s *d* = .10), as was the modulation of the (non-significant) response to pictures in the APC (Figure 1C, *t*(46) = 1.94, *p* = .06, Cohen’s *d* = 0.28). Null results cannot be interpreted as absence of effects, so we performed additional Bayesian one-sample *t*-tests to evaluate the support for the null model (absence of effect). These indicated moderate evidence for the null effect for odor association modulation in the PPC, both in the case of visual stimulation (B_01_ = 4.1) and auditory stimulation (B_01_ = 4.9). Similarly, there was weak evidence for the null model of odor association modulation in the APC for pictures (B_01_ = 1.1) and moderate support for the null model for sounds (B_01_ = 5.2). Details of the Bayesian t-tests can be found in the Supplementary material (Section 2.1).

Although we did not detect any effect of odor association on the activations in piriform cortex when considering the parametric modulations, a subset of objects with strong odor associations could potentially have produced an odor association-dependent activity in a non-linear fashion. To further investigate this possibility, we performed two additional analyses based on new GLMs. In the first analysis, we split the pictures and sounds into two groups, based on each individual’s odor association ratings, where one group comprised the lowest and one group the highest rated half of the stimuli. In the PPC ROI, we found that the visual and auditory activations persisted both for objects with weak and strong odor associations (weak: pictures with odor association 11.8 ± 10.9: *t*(40) = 4.19, *p* = .00015, Cohen’s *d* = 0.65; sounds with odor association 12.3 ± 11.7: *t*(40) = 7.52, *p* = 3.6*10^-9^, Cohen’s *d* = 1.17; strong: pictures with odor association 71.3 ± 16.1: *t*(40) = 3.79, *p* = .00050, Cohen’s *d* = 0.59; sounds with odor association 68.8 ± 16.1: *t*(40) = 7.68, *p* = 2.2*10^-9^, Cohen’s *d* = 1.20). Further, a Bayesian paired *t*-test showed moderate evidence for the null effect of no difference between these activations for both pictures (B_01_ = 5.26) and sounds (B_01_ = 5.89, see Supplementary materials Section 2.2 for details).

In the APC ROI, the auditory activation also remained significant for both objects with weak and strong odor associations (weak: *t*(40) = 6.48, *p* = 1.0*10^-7^, Cohen’s *d* = 1.01; strong: *t*(40) = 6.51, *p* = 9.2*10^-8^, Cohen’s *d* = 1.02). Again, a Bayesian paired *t*-test showed moderate evidence for the null effect of no difference between these activations (B_01_ = 5.61). The non-significant response to sounds in the APC seen in the main analysis remained non-significant both for objects with weak and strong odor associations (weak: *t*(40) = .42, *p* = 0.67, Cohen’s *d* = 0.07; strong: *t*(40) = 1.43, *p* = 0.16, Cohen’s *d* = 0.22), and the Bayesian paired *t*-test showed moderate evidence for the null effect of no difference between these responses (B_01_ = 4.53).

In the second analysis, we treated each stimulus as a separate condition and assessed the respective PPC ROI activations. We then ranked the stimuli according to each individual’s odor association ratings. These rankings did not significantly correlate with the activation for either pictures or sounds in the PPC (pictures: Spearman’s ρ = .19, *p* = .20, sounds: Spearman’s *ρ* = -.039, *p* = .79; Figure S2). In the APC, there was a significant correlation for pictures (Spearman’s *ρ* = .42, *p* = .0033), but not for sounds (Spearman’s *ρ* = -.073, *p* = .62). For comparison, we show the relationship between the activations and odor association ratings also for the auditory and visual ROIs in Figure S2.

Piriform cortex is not activated by nasal breathing but can be activated by sniffing alone, even in the absence of other sensory stimuli [Sobel et al., 1998]. Although participants were instructed to breathe normally through their nose and were not asked to sniff when presented with a stimulus, a potential confound is that participants might have non-consciously sniffed when presented with a visual or auditory stimulus [Perl et al., 2019], independent of its odor association. As a final control analysis, we therefore assessed whether the piriform activation was associated with a sniff response synchronized to stimulus onset; in other words, whether seeing or hearing a stimulus made participants initiate a sniff. To this end, we used the nasal respiratory data collected throughout the experiment and operationalized a “sniff” as an inhalation peak occurring at some point 0-1s after stimulus onset; a definition that biases our analyses towards finding sniff synchronization. This operationalization was used since the presented stimuli were not synchronized with breathing or presented with an inhalation cue. We then assessed the probability, considering the total number of nasal breaths throughout the recording, of a sniff occurring 0-1s after stimulus presentation. There was no significant increase in nasal inspiration time-locked to stimulus onset to either visual or auditory stimuli. Across all participants, the maximum probability of a peak occurring after a visual stimulus was .047 and occurring after a sound stimulus .050.

### Visual cortex processes auditory unisensory stimuli

We found cross effects in the two ROIs covering visual brain regions: Sounds activated V1 (*t*(46) = 8.01, *p* = 2.8*10^-10^, Cohen’s *d* = 1.17) and deactivated LOC (*t*(46) = -6.44, *p* = 6.4*10^-8^, Cohen’s *d* =-0.94; Figure 4A-B). Naturally, the sounds also activated multiple known nodes of the auditory processing network in the temporal lobe and frontotemporal junction (Figure S3). These expected activations not only indicate that our stimulation and analysis were effective but also serve as a data sanity check. Accordingly, the activations were significant in both A1 (*t*(46) = 21.76, *p* = 7.4*10^-26^, Cohen’s *d* = 3.17) and hAC (*t*(46) = 18.97, *p* = 2.2*10^-23^, Cohen’s *d* = 2.77; Figure 4D-E). In A1, the response to sounds was positively modulated by odor association (*t*(46) = 2.01, *p* = .0499, Cohen’s *d* = 0.29), whereas the modulation of the response to pictures was non-significant (*t*(46) = -0.13, *p* = .90, Cohen’s *d* = -0.02; Figure 4F). In hAC, none of the parametric modulations were significant (pictures: *t*(46) = -0.26, *p* = .79, Cohen’s *d* = -0.04; sounds: *t*(46) = 0.36, *p* = 0.72, Cohen’s *d* = 0.05).

**Figure 4.**
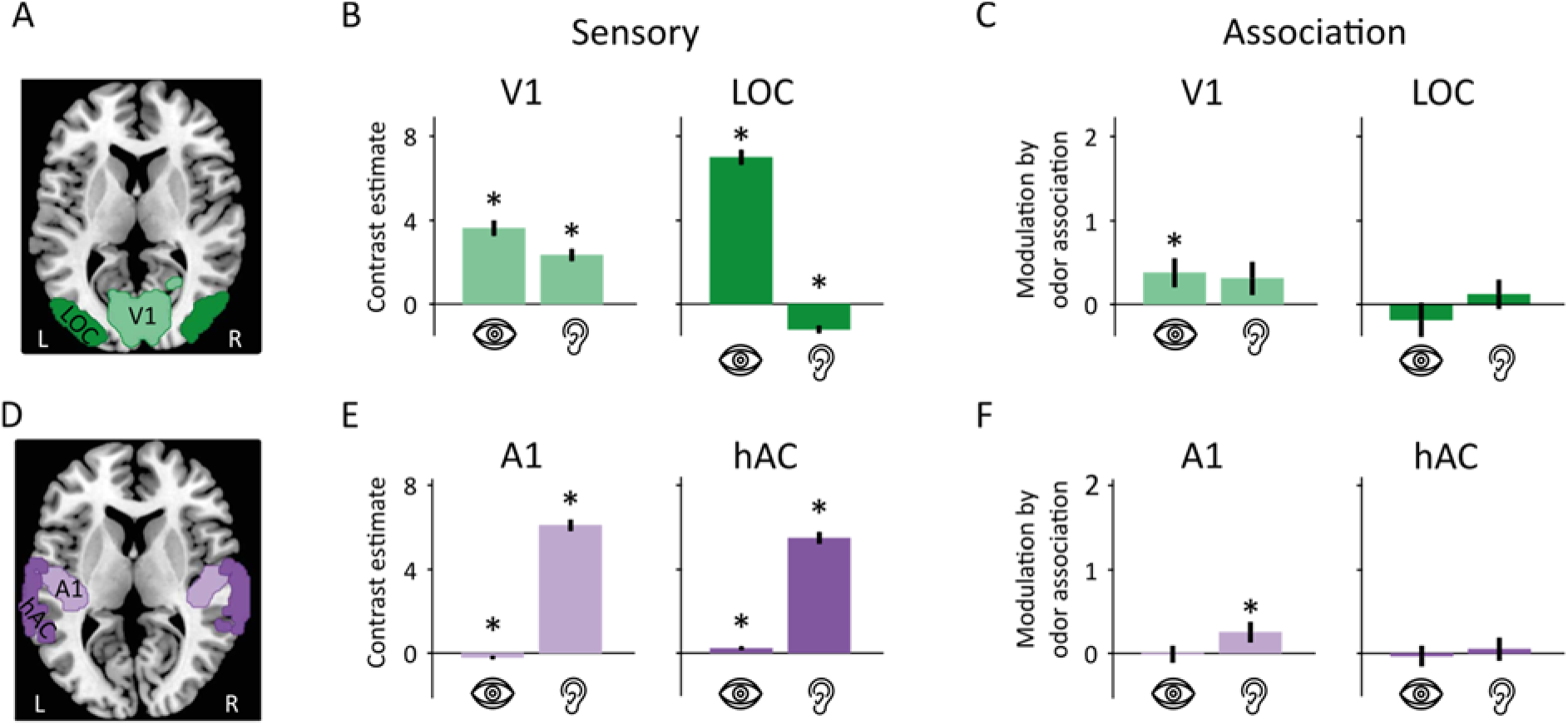
Region of interest MR imaging results for visual and auditory brain regions. **A**) Location of visual ROIs. B) Both sounds and pictures activated primary visual cortex (V1). In lateral occipital complex (LOC), activity increased in response to pictures and decreased in response to sounds. **C**) In V1, the modulation by odor association was significantly above zero for pictures, but not for sounds. In LOC, there was no significant modulation for either pictures or sounds. **D**) Location of auditory ROIs. **E**) In primary auditory cortex (A1), the BOLD signal increased in response to sounds and decreased in response to pictures. Both pictures and sounds elicited significant activations in higher order auditory cortex (hAC). **F**) In A1, the modulation by rated odor association was significantly above zero for sounds, but not for pictures. Asterisks mark statistical significance and error bars depict ± 1 Standard Error of the Mean (SEM). Note the varying scale on the y axes.

### Auditory cortex processes visual unisensory stimuli

Cross-modal effects were also noted in the auditory ROIs: Pictures elicited a decreased neural activity in primary auditory cortex (A1; *t*(46) = -2.45, *p* = .018, Cohen’s *d* = -0.36) and an increased activity in higher auditory cortices (hAC; *t*(46) = 2.42, *p* = .019, Cohen’s *d* =0.35; Figure 4E). Naturally, the visual stimulation also activated multiple regions in the occipital and parietal cortices that are all known nodes in the visual processing network (Figure S3), again indicating that our stimulation and analysis worked as intended. These activations were significant both in primary visual cortex (V1; *t*(46) =9.79, *p* = 7.97*10^-13^, Cohen’s *d* = 1.43) and in the visual object-processing area LOC (*t*(46) = 8.01, *p* = 2.83*10^-10^, Cohen’s *d* = 1.17; Figure 4B). In V1, the activation by pictures was positively modulated by odor association (*t*(46) = 2.16, *p* = .036, Cohen’s *d* = 0.32), whereas the activation by sounds was not (*t*(46) = 1.53, *p* = .13, Cohen’s *d* = 0.22; Figure 4C). In LOC, none of the parametric modulations were significant (pictures: *t*(46) = -0.89, *p* = .38, Cohen’s *d* = -0.13; sounds: (*t*(46) = 0.68, *p* = .50, Cohen’s *d* = 0.10).

### Transfer of auditory and visual object information through the amygdala to the PPC

We demonstrated above that PPC processes unisensory visual and auditory objects. There are, however, no known monosynaptic connections between PPC and either the visual or auditory cortices. Hence, we next sought to understand which cerebral node might mediate the information to the PPC. To this end, we assessed the condition relevant shortest path between the auditory and visual object-oriented cortices and PPC based on dynamic functional connectivity dFC). We found that both auditory and visual object areas convey functional information to the PPC via the amygdala [x ± 26.25, y -8.25, z -26.25]; Pictures: LOC--AMY *t*(46) = 17, *p* < .0001, AMY—PPC, *t*(46) = 17, *p* < .0001; Sound: hAC—AMY, *t*(46) = 13, *p* < .0001, AMY--PPC, *t*(46) = 17, *p* < .0001 (Figure 5E).

**Figure 5.**
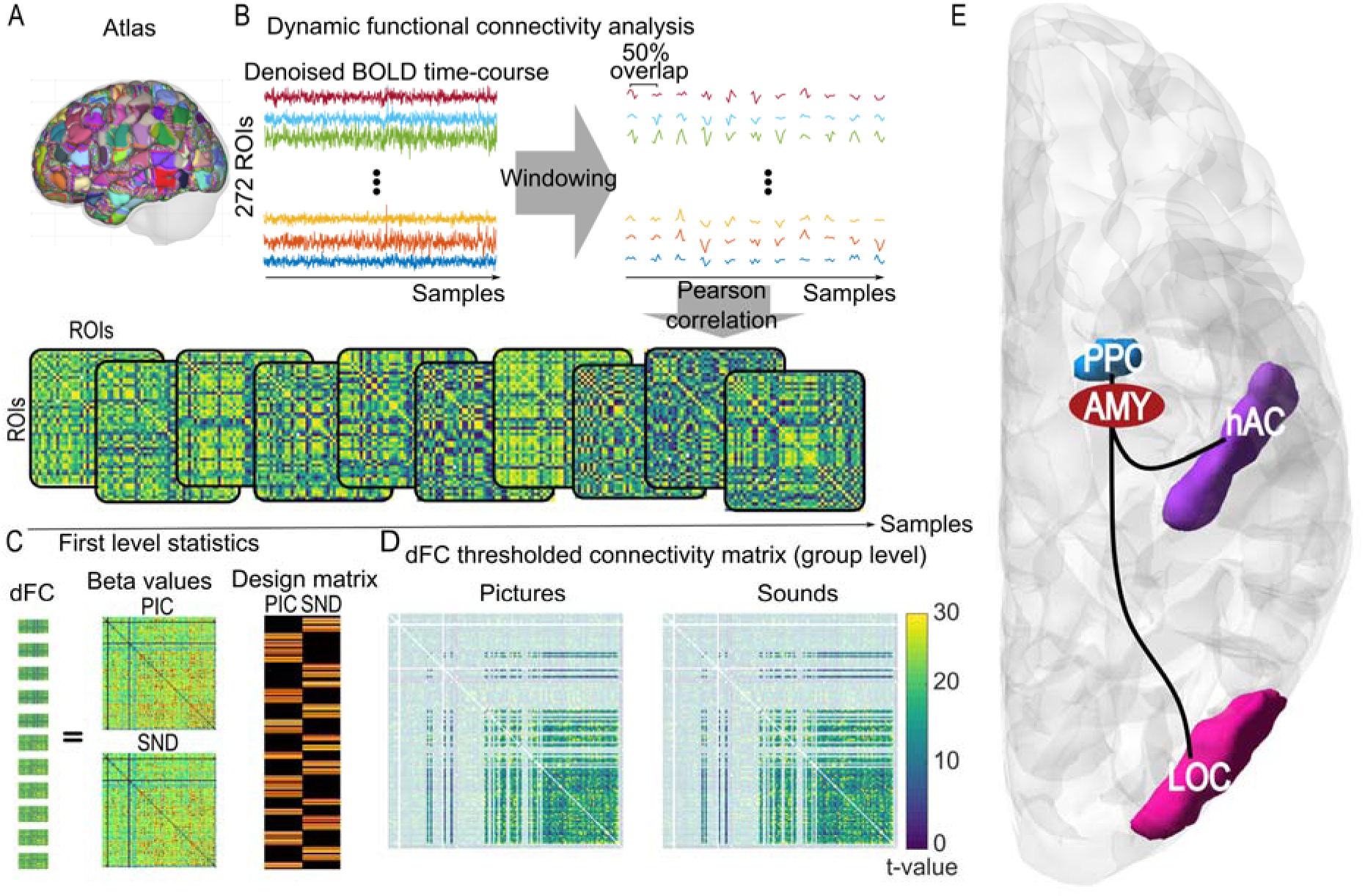
The dFC analysis determined the condition-relevant shortest functional path between object-oriented cortices and PPC. **A**) The 272 ROI atlas covering the whole brain. **B**) Denoised and windowed BOLD signals were extracted from the 272 ROIs. Pearson correlation was used to construct the functional connectivity for each window, cumulatively creating brain-wide dynamic functional connectivity (dFC). **C**) Linear regression of stimulus-dependent modulation of the dFC at the individual level. The design matrix indicates samples relevant to each stimulus (i.e., sounds and pictures). Effects were estimated as individual Beta values. **D**) Group level Beta values were *t*-transformed and thresholded based on the average gray matter of ROIs in the cohort. The small brain map in the middle indicates the averaged gray matter probability that was used for the thresholding of the group level t-transformed dFCs. **E**) Finally, we calculated the condition-relevant shortest path between the audio (hAC) and visual (LOC) object-oriented cortices and PPC.

## DISCUSSION

We here aimed to distinguish between two main candidate explanations of unisensory activation of the olfactory cortex by non-olfactory stimuli: activation by the non-olfactory stimulus per se (true unisensory processing of non-preferred stimuli), and activation due to olfactory association to the stimulus (cross-modal association). We first demonstrated that piriform (olfactory) cortex responded to both unimodal visual and unimodal auditory stimuli in form of pictures and sounds of object which were neither explicitly nor implicitly linked to odors during the experiment. Specifically, we found that the posterior piriform cortex (PPC), an area known to process odor objects [Gottfried, 2010; Howard et al., 2009], responded to both pictures and sounds of objects. The anterior piriform cortex (APC), which is thought to code mainly for low-level odor properties [Gottfried, 2010], responded to sounds of objects. Thereafter, we demonstrated that the piriform activation by non-olfactory unisensory stimuli did not depend on either the odor association of the presented objects or a sniff response to their presentation. Together, these results provide support for true unisensory non-olfactory processing in olfactory cortex.

Our results demonstrate that unisensory auditory and visual objects can activate piriform cortex, even in the absence of any odor stimulation or mention of odors. Critically, this observed cross-modal processing in piriform cortex was independent of the individual’s level of odor association to the different objects (rated post fMRI acquisition), as demonstrated by three independent analyses. First, there was moderate support for the null model of an absence of modulation by odor association. Second, there was clear activation of piriform cortex even for objects that the participant associated only weakly with odors, which was also comparable to the effects for objects with strong odor associations. Third, we found no correlation between odor association and piriform activation. Our results extend prior findings of cross-modal processing in human piriform cortex [Gottfried et al., 2004; Karunanayaka et al., 2015; Porada et al., 2019; Schulze et al., 2017] in two important ways. We show both auditory and visual cross-modal activity in the piriform cortex whereas previous studies have only reported visual cross-modal activity. A recent study by Porada and colleagues [2019] included unisensory auditory stimulation but did not find auditory evoked activation of piriform cortex; possibly due to a lack of sufficient statistical power to detect small effects. The present study has considerably higher statistical power with three times as many participants and roughly five times as many trials per participant. Second, and more importantly, all previous studies associated the presented non-olfactory stimuli with olfactory stimuli before and/or during neuroimaging data collection. For example, both Porada and colleagues [2019] and Gottfried and colleagues [2004] experiments were preceded by a session where the associations between the non-olfactory and olfactory stimuli were learned. The non-olfactory stimuli giving rise to piriform activity were thus explicitly associated with odors and it is likely that participants associated all stimulation to odors, even when no odor was simultaneously presented. We did not present or mention odors prior to or during fMRI data acquisition, making it possible to distinguish between true sensory processing of non-preferred stimuli and cross-modal association. Because we did not want to prompt participants to think about odors during the experiment, we did not use any functional olfactory region localizer. Such a localizer would perhaps be more precise, but it would require explaining to participants, prior to the experiment, that they would be stimulated with odors. Therefore, we instead used the same functionally and anatomically constrained ROIs as used in Porada (2019) to preserve independence between ROI and data analyses.

We have clearly demonstrated that the cross-modal processing of auditory and visual stimuli in piriform cortex is not dependent on how much the participant associates the presented object with an odor. That said, the cross-modal activity may still depend on the fact that our unimodal auditory and visual stimulation depicted objects. The activation was significant for both auditory and visual object stimuli in the PPC. Given that this region is commonly linked to olfactory object and category processing [Gottfried, 2010; Howard et al., 2009], it is plausible that the processing of non-olfactory input should be equally dependent on object or category specific content. This hypothesis is supported by our recent study demonstrating multisensory integration of object information in PPC [Porada et al., 2019]. Specifically, olfactory, auditory, and visual object stimuli were presented uni-, bi-, and trimodally and the activity in PPC increased with the number of sensory modalities providing object information, but only when the object information was congruent (i.e., the same object presented) in the different sensory modalities. This indicates that specific object-related information, rather than low-level stimulus properties alone, affects the activity in PPC also in the case of non-olfactory stimuli. We propose that this non-olfactory object processing is what we observe in PPC in the current study. We furthermore speculate that the auditory activation of APC might be caused by the relatively high auditory activation of PPC as a reflection of the strong internal connectivity between the auditory cortex and the PPC [Carmichael et al., 1994]. Based on the current data, we cannot conclusively determine whether the piriform activity evoked by pictures and sounds of objects indeed reflects processing of object and not low-level information. Future studies could address this by either contrast the evoked activity from auditory and visual object and non-object stimuli with similar low-level perceptual properties or, conversely, determine whether visual/auditory objects can be decoded based on piriform cortex activation patters.

An alternative interpretation of the current findings is that the cross-modal processing in piriform cortex is not related to the specific object information or low-level properties of the stimuli, but instead is an effect caused by increased attention or arousal evoked by the onset of the auditory and visual stimuli. Indeed, attention-dependent activity in piriform cortex has been demonstrated, but only in the APC [Zelano et al., 2005], a region where we in the current study find cross-modal activation only by auditory stimuli. It should, however, also be noted that the attention effect demonstrated by Zelano and colleagues [2005] was evident when attending to odors, whereas the participants in the current study were unaware that the study in fact assessed phenomena related to odors and were presumably not engaged in any conscious odor-searching behavior. Speaking against the notion that our obtained results are mediated by attentional processes is further the fact that we did not find a stimuli-synchronized sniff response, a phenomenon argued to be associated with attentional allocation [Perl et al., 2019]. Nonetheless, it is possible that our findings are generated by a general preparatory or prediction coding effect where non-olfactory sensory input prepares olfactory cortex for potential incoming olfactory input in response to new non-olfactory information [Ohla et al., 2017].

The functional connectivity path analysis provides us with potential paths that the auditory and visual object information travels from object processing areas in the auditory (hAC) and visual cortex (LOC) to the olfactory object processing region (PPC). Both the visual and auditory related information is seemingly conveyed from their respective cortices to the PPC via the amygdala. The amygdala is a well-known, and integral, part of the general olfactory system via its monosynaptic connection to both the olfactory bulb and the piriform cortex with especially strong connection to the posterior piriform cortex [Mainland et al., 2014; Majak et al., 2004]. Amygdala is also uniquely situated to convey information from both the visual and auditory system via its cortical inputs from their respective cortical sensory systems [Mosher et al., 2010] as well as its rich multisensory neuron population [Shan et al., 2023]. To date, it is not clear what role the amygdala might serve within the olfactory system but as noted by Noto and colleagues [2021], the amygdala might be involved in reward processing and the generation of rapid approach/avoid responses. One can therefore speculate whether the function of the visual/auditory activation of the piriform cortex is to subserve the olfactory system’s task of assisting us to make rapid approach/avoidance decisions [Arshamian et al., 2017; Iravani et al., 2021].

Analogously to the crossmodal activity of the olfactory cortex, we found cross-modal effects also in the visual and auditory ROIs. Primary visual cortex (V1) was activated by the sounds of objects, and hAC was activated by the pictures. It is here interesting to note that the latter of these cross-activations was apparent also in the aforementioned study by Porada and colleagues [2019]. In primary auditory cortex (A1) and LOC, we instead found decreased BOLD signal in response to the non-preferred stimulation (pictures and sounds, respectively). Similar effects have previously been described in auditory and extrastriate visual areas [Laurienti et al., 2002]. Moreover, there are several other accounts of cross-modal effects in the auditory and visual cortices on a finer spatial scale. For example, over 40% of cells in the cat visual parastriate cortex responds to unimodal localized auditory stimuli [Morrell, 1972], auditory stimulation influences the preferences of cells in the cat V1 area [Chanauria et al., 2019], and visual and auditory objects can be decoded from auditory and visual cortex, respectively, in humans [Hsieh et al., 2012; Meyer et al., 2010; Vetter et al., 2014]. Thus, previous work has demonstrated stimulus-specific activation in auditory cortex by unimodal visual stimulation, and stimulus-specific activation in visual cortex by unimodal auditory stimulation. We can here conceptually replicate the general finding of visual-auditory cross-activation.

It has previously been argued that the neocortex, including both higher-order processing regions and regions traditionally thought of as unisensory, is in essence multisensory [Ghazanfar and Schroeder, 2006]. We can here provide support to this notion and extend this concept of a multisensory neocortex by demonstrating that the olfactory cortex, which is evolutionarily the oldest sensory cortex and not part of the neocortex which evolved from the olfactory-dominated cerebral hemispheres [Northcutt and Kaas, 1995], also appear to respond in a multisensory fashion: It appears to process non-olfactory input independently of odor association. Moreover, it has been proposed that the brain is organized according to tasks rather than sensory input, with the possible exception of primary sensory cortices [Amedi et al., 2017]. Studies on congenitally blind individuals have shown that they demonstrate the same cortical specialization in ‘visual’ cortex for non-visual stimuli as is normally seen in visual processing, such as intact ventral/dorsal pathways [Cecchetti et al., 2016]. Assuming that the cross-modal activity in the PPC that we have demonstrated reflects processing of non-olfactory object information, our findings are consistent with a task-oriented brain organization in which the role of PPC is object processing, with a preference for olfactory input. Another interpretation of our results is that the PPC is an olfactory region that can be activated by non-olfactory input in a preparatory way, possibly driven by a lifetime of multisensory input. Regardless of which theory will prevail, our findings support the notion of a multimodal cortex and highlights that this extends to the olfactory cortex.

While we did not explicitly test participants on the identification task, we are confident that they in general paid attention to and were able to identify the stimuli for several reasons. First, we monitored their eye-tracking and participants with excessive eye closure were excluded. Second, the strong visual and auditory activations in the major respective processing areas show that the stimuli were in general seen and heard. Third, the practice task allowed participants to familiarize themselves with the objects and learn their identities. Given that all objects are in our everyday life commonly occurring items, it is unlikely that participants were not able to identify all stimuli.

In summary, we here conclusively demonstrate true unisensory processing of sounds and pictures of objects in olfactory cortex that is independent of odor associations. These results provide additional support against the notion of piriform cortex as a true unisensory olfactory region and supports the argument that sensory processing is best characterized as relying on multisensory networks.

## Acknowledgments

Funding provided by the Knut and Alice Wallenberg Foundation (KAW 2018.0152) and the Swedish Research Council (2021-06527), awarded to JNL. Data acquisition was supported by a grant to the Stockholm University Brain Imaging Centre (SU FV-5.1.2-1035-15).

## Notes

https://osf.io/duwrb/?view_only=d214b458aee64204bf6e09df38249631

### Competing Interest Statement

The authors have declared no competing interest.

### Summary of Updates

Updated analyses and results of connectivity analyses.

