## Supplementary for "Unisensory visual and auditory objects are processed in olfactory cortex, independently of odor association"

### Supplementary material

#### 1. Pilot task

The pilot study on 23 healthy participants (14 female, mean age 28 years, SD 6.9 years) was conducted with the aim to identify a suitable stimulus set for the fMRI experiment. Based on the outcome of the pilot study, we selected objects for which both the sound and the image were easy to identify, and where the sound and image evoked a similar degree of odor association.

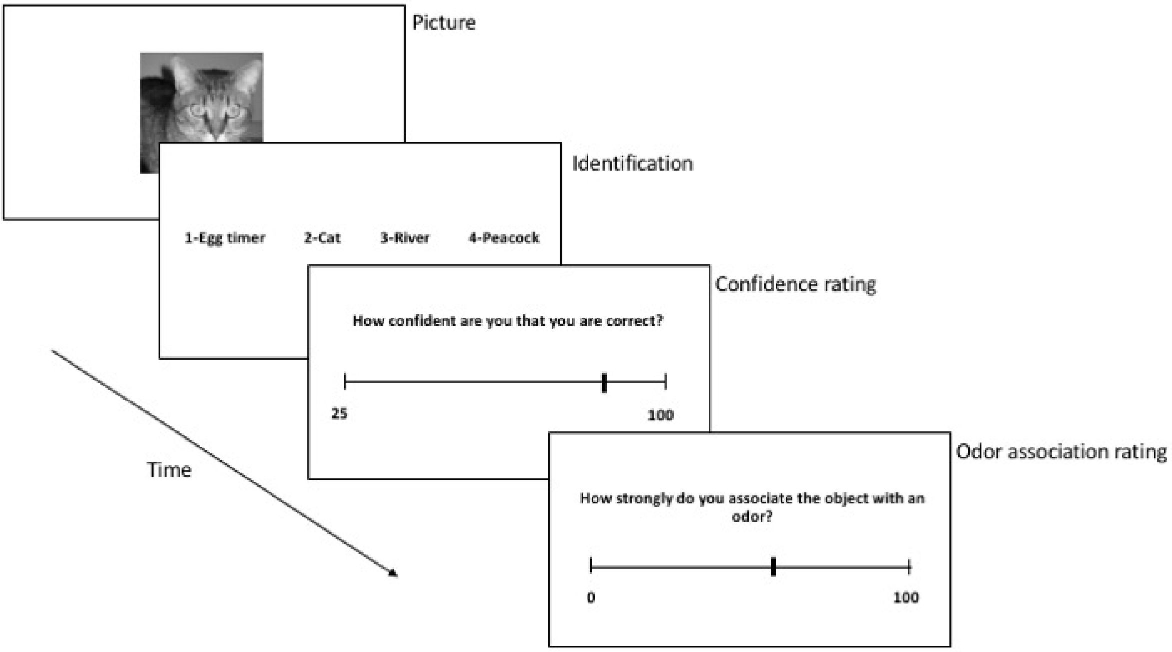


**Figure S1. Pilot study procedure.** Participants identified the object represented by each image and each sound (separately) in a 4-alternative forced-choice task where the four options were designed to include the correct answer, a similar but wrong answer, a randomly chosen object out of the 59 other objects, and an obviously wrong answer. In the next step, participants rated how confident they were of their response on a visual analogue scale ranging from 0 (no odor association) to 100 (very strong odor association. Last, participants rated to what extent they associated the object with an odor. An example trial with a picture (a cat) is depicted here.

#### 2. Bayesian tests

##### *2.1 Effects of odor association modulation*

Bayesian t-tests were performed in JASP, to evaluate the null effects of parametric modulation by the individual odor associations in the posterior piriform cortex (PPC) and anterior piriform cortex (APC) region of interests. The obtained results from these are presented below.

**PPC - visual stimulation:**

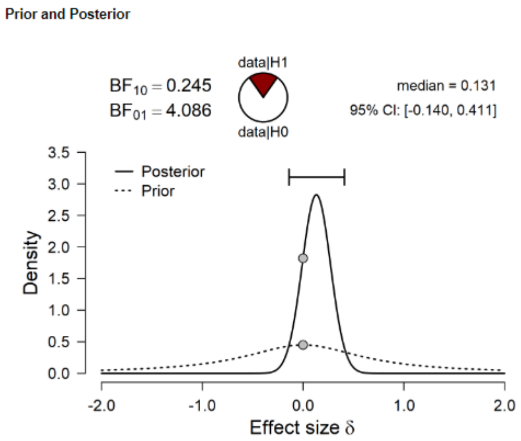


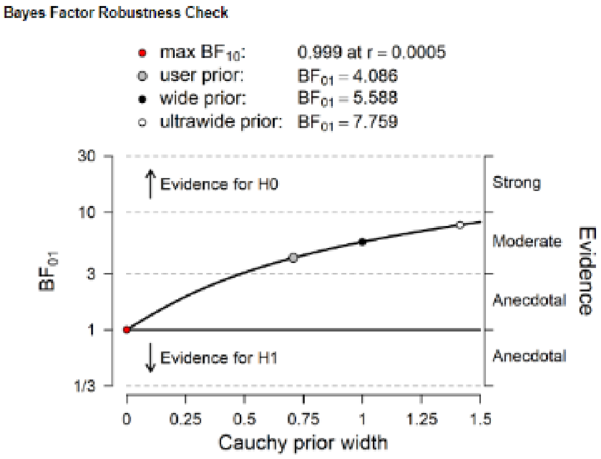


**PPC - auditory stimulation:**

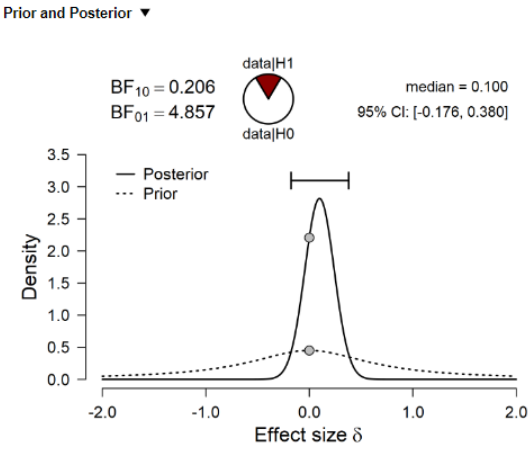


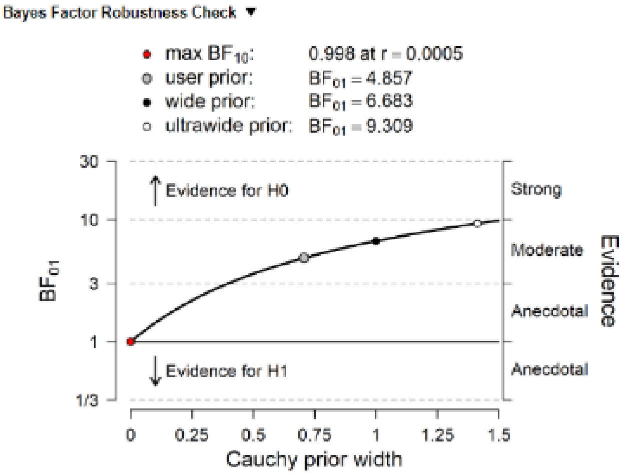

#### APC - visual stimulation:

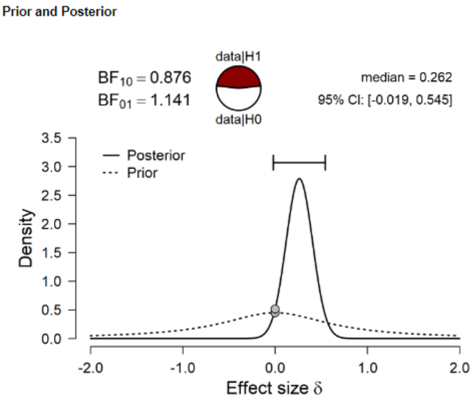


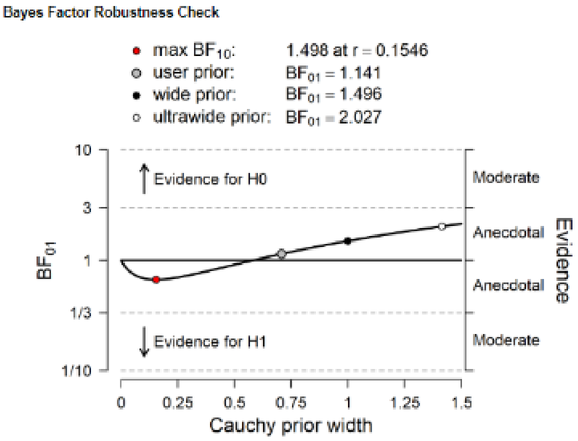


#### APC - auditory stimulation

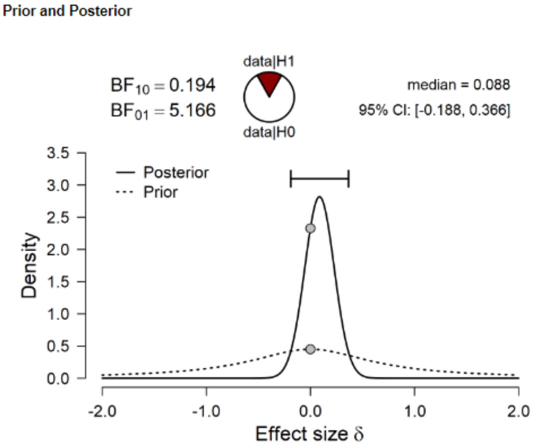


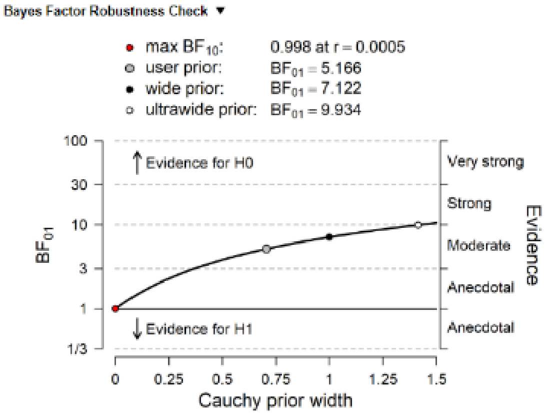

#### 2.2 Comparison of activations from objects with weak and strong odor association

##### PPC - visual stimulation: Weak versus strong odor association

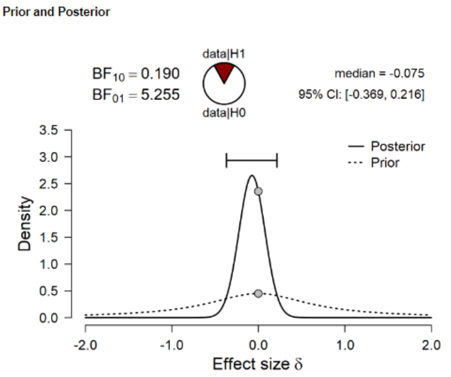


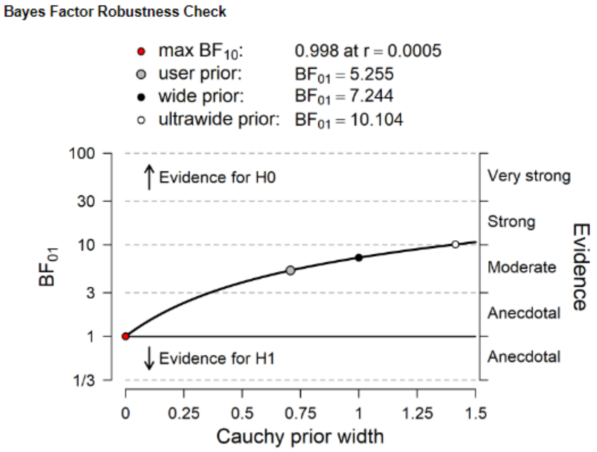


##### PPC - auditory stimulation: Weak versus strong odor association

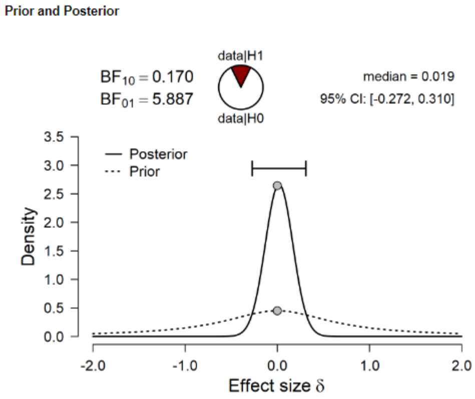


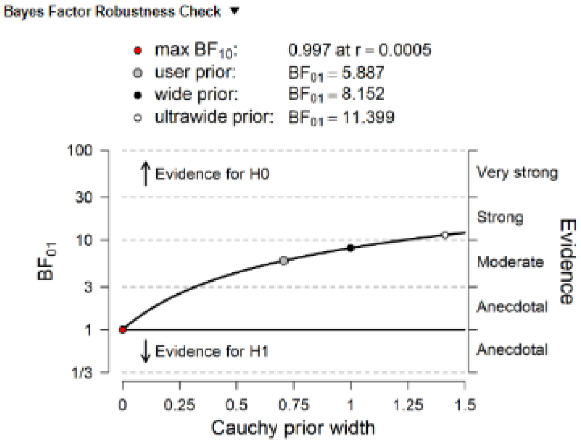

#### APC - visual stimulation: Weak versus strong odor association

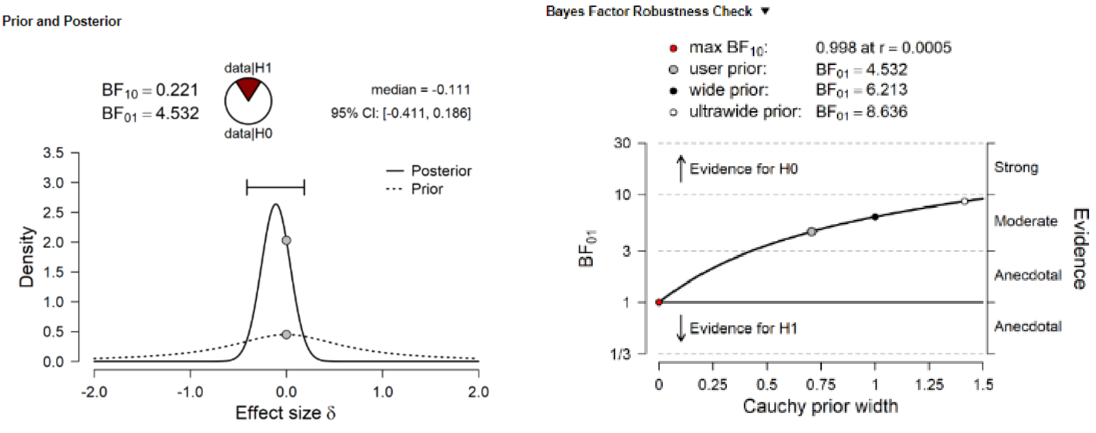


#### APC - auditory stimulation: Weak versus strong odor association

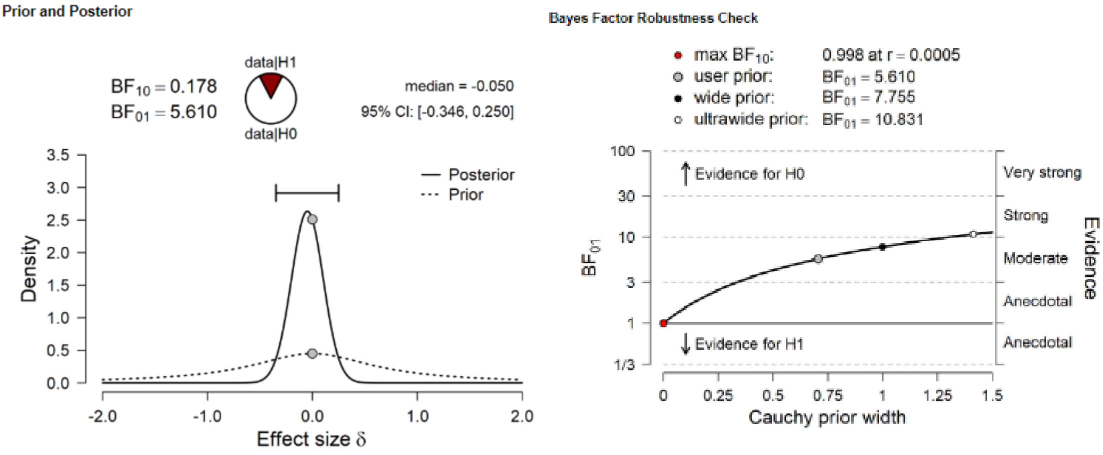

#### 3. Correlation with odor association

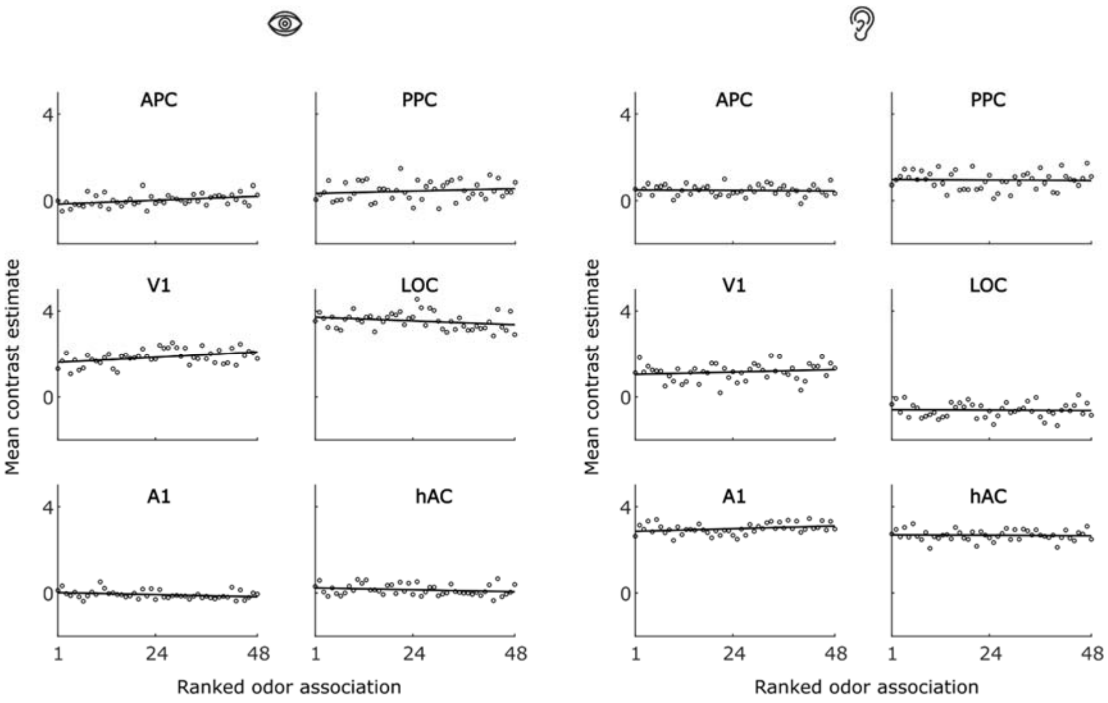


**Figure S2.** BOLD activations in the ROIs as a function of ranked odor association level for the pictures (left; eye symbol) and sounds (right; ear symbol) of the objects. The 48 objects were rank sorted based on their rated odor association, individually per participant and separately for pictures and sounds. The marks depict participant-average data. Solid black lines are best linear fits.

#### 4. Whole-brain visual and auditory activation

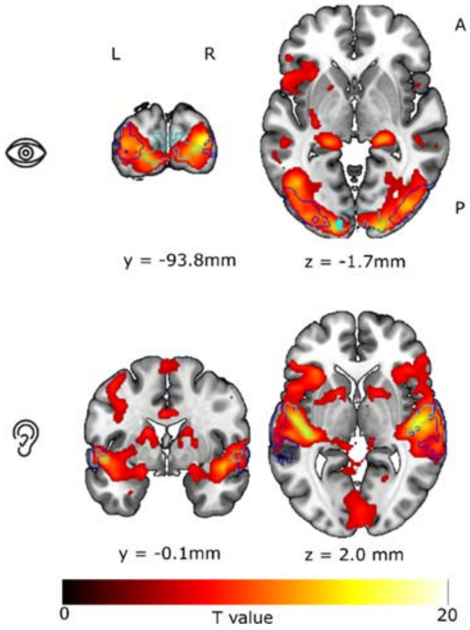


**Figure S3:** Whole-brain fMRI results overlaid on the MNI152 default template image in MRIcroGL. Upper panel: Visual activation. The outlines of the visual ROIs are overlaid in cyan (V1) and blue (LOC). Lower panel: Auditory activation. The outlines of the auditory ROIs are overlaid in cyan (A1) and blue (hAC). Both the visual and the auditory activations are corrected for multiple comparisons using a peak-level threshold at $p$ = .05 family-wise error (FWE) across the whole field of view.
